# Interactive Visual Analysis of Mass Cytometry Data by Hierarchical Stochastic Neighbor Embedding Reveals Rare Cell Types

**DOI:** 10.1101/169888

**Authors:** Vincent van Unen, Thomas Höllt, Nicola Pezzotti, Na Li, Marcel J. T. Reinders, Elmar Eisemann, Frits Koning, Anna Vilanova, Boudewijn P. F. Lelieveldt

**Affiliations:** Department of Immunohematology and Blood Transfusion, Leiden University Medical Center, Albinusdreef 2, 2333 ZA, Leiden, the Netherlands; Computer Graphics and Visualization Group, Mekelweg 4, 2628 CD, Delft University of Technology, Delft, the Netherlands; Computational Biology Center, Leiden University Medical Center, Albinusdreef 2, 2333 ZA, Leiden, the Netherlands; Pattern Recognition and Bioinformatics Group, Delft University of Technology, Mekelweg 4, 2628 CD, Delft, the Netherland; Division of Image Processing, Department of Radiology, Leiden University Medical Center, Albinusdreef 2, 2333 ZA, Leiden, the Netherlands

**Keywords:** Mass cytometry, t-Distributed Stochastic Neighbor Embedding, t-SNE, Hierarchical SNE, HSNE, rare cells, single-cell analysis, high-dimensional data analysis

## Abstract

Mass cytometry allows high-resolution dissection of the cellular composition of the immune system. However, the high-dimensionality, large size, and non-linear structure of the data poses considerable challenges for data analysis. In particular, dimensionality reduction-based techniques like t-SNE offer single-cell resolution but are limited in the number of cells that can be analysed. Here we introduce Hierarchical Stochastic Neighbor Embedding (HSNE) for the analysis of mass cytometry datasets. HSNE constructs a hierarchy of non-linear similarities that can be interactively explored with a stepwise increase in detail up to the single-cell level. We applied HSNE to a study on gastrointestinal disorders and three other available mass cytometry datasets. We found that HSNE efficiently replicates previous observations and identifies rare cell populations that were previously missed due to downsampling. Thus, HSNE removes the scalability limit of conventional t-SNE analysis, a feature that makes it highly suitable for the analysis of massive high-dimensional datasets.

## Introduction

Mass cytometry (cytometry by time-of-flight; CyTOF) allows the simultaneous analysis of multiple cellular markers (>30) present on biological samples consisting of millions of cells. Computational tools for the analysis of such datasets can be divided into clustering-based and dimensionality reduction-based techniques^1^, each having distinctive advantages and disadvantages. The clustering-based techniques, including SPADE^2^, FlowMaps^3^, Phenograph^4^, VorteX^5^ and Scaffold maps^6^, allow the analysis of datasets consisting of millions of cells but only provide aggregate information on generated cell clusters at the expense of local data structure (i.e. single-cell resolution). Dimensionality reduction-based techniques, such as PCA^7^, t-SNE^8^ (implemented in viSNE^9^), and Diffusion maps^10^, do allow analysis at the single-cell level. However, the linear nature of PCA renders it unsuitable to dissect the non-linear relationships in mass cytometry data, while the non-linear methods (t-SNE^8^ and Diffusion maps^10^) do retain local data structure, but are limited by the number of cells that can be analyzed. This limit is imposed by a computational burden but, more importantly, by local neighborhoods becoming too crowded in the high-dimensional space resulting in overplotting and presenting misleading information in the visualization. In cytometry studies this poses a problem, as a significant number of cells needs to be removed by random downsampling to make dimensionality reduction computationally feasible and reliable. Future increases in acquisition rate and dimensionality in mass- and flow cytometry are expected to amplify this problem significantly^11,12^.

Here, we adapted Hierarchical Stochastic Neighbor Embedding (HSNE)^13^ that was recently introduced for the analysis of hyperspectral satellite imaging data to the analysis of mass cytometry datasets to visually explore millions of cells while avoiding downsampling. HSNE builds a hierarchical representation of the complete data that preserves the non-linear high-dimensional relationships between cells. We implemented HSNE in an integrated single-cell analysis framework called Cytosplore^+HSNE^ This framework allows interactive exploration of the hierarchy by a set of *embeddings*, two-dimensional scatter plots where cells are positioned based on the similarity of all marker expressions simultaneously, and used for subsequent analysis, such as clustering of cells at different levels of the hierarchy. We found that Cytosplore^+HSNE^ replicates the previously identified hierarchy in immune-system-wide single-cell data^4,5,14^, i.e. we can immediately identify major lineages at the highest overview level, while acquiring more information by dissecting the immune system at the deeper levels of the hierarchy on demand. Additionally, Cytosplore^+HSNE^ does so in a fraction of the time required by other analysis tools. Furthermore, we identified rare cell populations specifically associating to diseases in both the innate and adaptive immune compartments that were previously missed due to downsampling. We highlight scalability and generalizability of Cytosplore^+HSNE^ using two other datasets, consisting of up to 15 million cells. Thus, Cytosplore^+HSNE^ combines the scalability of clustering-based methods with the local single-cell detail preservation of non-linear dimensionality reduction-based methods. Finally, Cytosplore^+HSNE^ is not only applicable to mass cytometry datasets, but can be used for other high-dimensional data like single-cell transcriptomic datasets.

## Results

### Hierarchical Exploration of Massive Single-Cell Data

For a given high-dimensional dataset such as the three-dimensional illustrative example in Figure 1a, HSNE^13^ builds a hierarchy of local neighborhoods in this high-dimensional space, starting with the raw data that, subsequently, is aggregated at more abstract hierarchical levels. The hierarchy is then explored in reverse order, by embedding the neighborhoods using the similarity-based embedding technique, Barnes-Hut (BH)-SNE^15^. To allow for more detail and faster computation, each level can be partitioned in part or completely, by manual gating or unsupervised clustering, and partitions are embedded separately on the next, more detailed level (compare Fig. 1b). HSNE works particularly well for the analysis of mass cytometry data, because the local neighborhood information of the data level is propagated through the complete hierarchy. Groups of cells that are close in the Euclidian sense (Fig. 1a, grey arrow), but not on the non-linear manifold (Fig. 1a, dashed black line), are well separated even at higher aggregation levels (Fig. 1b). The power of HSNE lies in its scalability to tens of millions of cells, while the possibility to continuously explore the hierarchy allows the identification of rare cell populations at the more detailed levels. Next follows a general description of how the hierarchy is built and explored through embeddings. More details can be found in the **Methods**.

**Figure 1.**
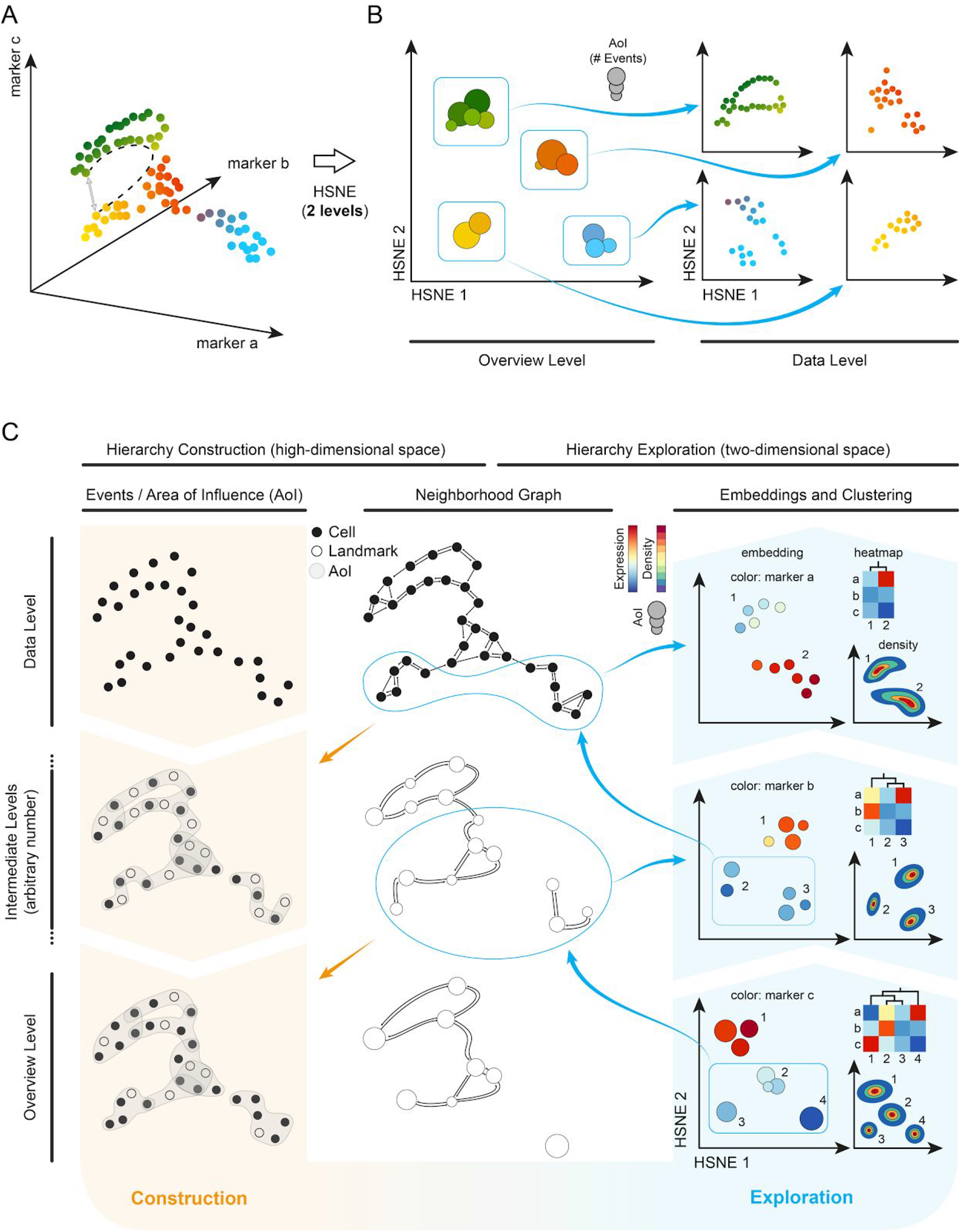
Schematic overview of Cytosplore^+HSNE^ for exploring mass cytometry data. By creating a multi-level hierarchy of an illustrative 3D dataset **(a)** we achieve a clear separation of different cell groups in an overview embedding (left panel **b**) that conserves non-linear relationships (i.e. follows the distance indicated by the dashed line in panel a instead of the grey arrow) and more detail within the separate groups on the data level (right panel **b**). **(c)** Construction and exploration of the hierarchy. The hierarchy is constructed starting with the data level (left two columns). Based on the high-dimensional expression patterns of the cells a weighted kNN graph is constructed, which is used to find representative cells used as landmarks in the next coarser level. By administering the area of influence (AoI) of the landmarks, cells/landmarks can be aggregated without losing the global structure of the underlying data or creating shortcuts. The exploration of the hierarchy is shown in the two rightmost columns. At the bottom we see the overview level (in this example the 3^rd^ level in the hierarchy), which shows that a group of landmarks has low expression in marker c (bottom-right panel). Selecting this group of landmarks for further exploration results in a look-up of the landmarks in the preceding level (neighborhood graph, intermediate level) that are in the AoI, with which a new embedding can be created at the 2^nd^ level of the hierarchy (middle-right panel). Marker b shows a strong separation between the upper and lower landmarks at this level. Zooming-in on the landmarks with low expression of marker b reveals further separation in marker a at the lowest level, the full data level (top-right panel).

#### Hierarchy Construction

The left panels of Figure 1c give an overview of the HSNE-hierarchy construction. We show the hierarchy from the fine-grained data level to an overview level from the top to bottom panels. The number of levels is defined by the user and depends mostly on the input-data size. We recommend to use log10(N/100) levels, with N being the number of cells. The foundation of the hierarchy is constructed using the original input data. Each dot represents a single cell (Fig. 1c, data level). Similarities between cells on the data level are defined by building an approximated, weighted k-nearest neighbor (kNN) graph^16^ (Fig. 1c, top-center panel). The weights of this graph can directly be used as input to embed the data into a two-dimensional space (Fig. 1c, top-right panel). With the BH-SNE the two-dimensional embedding is generated such that the layout of the points indicates similarities between the cells in the high-dimensional space according to the neighborhood graph.

#### Aggregation of data

To aggregate the data into the next level (Fig. 1c, intermediate levels), we identify representative cells to use as landmarks (Fig. 1c, white circles). For that, the weighted kNN graph is interpreted as a Finite Markov Chain and the most influential (i.e., best-connected) nodes are chosen as landmarks, using a Monte Carlo process. The landmarks are then embedded into a two-dimensional space based on their similarities. However, simply repeating the kNN construction for the selected landmarks in the high-dimensional space would eventually eliminate non-linear structures by creating undesired “shortcuts” in the graph (a problem reported by Setty et al.^17^ in a different setting). Instead, we define the area of influence (Aol) of each landmark, indicated by the grey hulls (Fig. 1c, left panels), as the cells that are well-represented by the landmark according to the kNN graph. Different landmarks can have overlapping regions of locally-similar cells. Therefore, we define the similarity of two landmarks as the overlap of their respective Aols. Furthermore, we construct a neighborhood graph, based on these similarities, that replaces the kNN graph as input for levels subsequent to the data level. Hereby, we effectively maintain the non-linear structure of the data to the top of the hierarchy and avoid shortcuts (Fig. 1c, bottom panels). We show that the preservation of non-linear neighborhoods by HSNE indeed conserves structure that is otherwise lost by random downsampling (Supplementary Notes 1 and Supplementary Fig. 1).

#### Interactive Exploration

Data exploration in Cytosplore^+HSNE^ starts with the visualization of the embedding at the highest level, the overview level (Fig. 1c, bottom-right panel). Similar to other embedding techniques for visualizing single-cell data^4,9^, the layout of the landmarks indicates similarity in the high-dimensional space according to the level’s neighborhood graph. Color is used to represent additional traits, such as marker expressions. The landmark size reflects its AoI. While it is possible to continuously select all landmarks and compute a complete embedding of the next, more detailed level, this strategy would eventually embed all data and suffer from the same scalability problems as a t-SNE embedding, i.e., overcrowding (Supplementary Notes 2 and Supplementary Fig. 2) and slow performance. Instead, we envision that the user selects a group of landmarks, by manual gating based on visual cues, such as patterns found in marker expression, or by performing unsupervised Gaussian Mean Shift (GMS) clustering^18^ of the landmarks based on the density representation of the embedding (Fig. 1c, right panels). Then, the user can zoom into this selection by means of a more detailed embedding that consists of the landmarks/cells in the combined Aol on the preceding level. Moreover, interactively linked heatmap visualizations of clusters (Fig. 1c, right panels) and descriptive statistics of markers within a selection can be used to guide the exploration. Importantly, all of the described tools are available at every level of the hierarchy and linked interactively. Selections in the embedding and heatmap at one level of the hierarchy can thus be highlighted in the embeddings of other levels (Supplementary Fig. 3). All these aspects are further demonstrated using a typical exploration workflow with Cytosplore^+HSNE^ in the Supplementary Video 1. With this strategy, tens of millions of cells can be explored, providing both global visualizations up to single-cell resolution visualizations, while preserving non-linear relationships between landmarks/cells at all levels of the hierarchy.

### Cytosplore^+HSNE^ Eliminates the Need for Downsampling

In a previous study^14^, a mass cytometry dataset on 5.2 million cells derived from intestinal biopsies and paired blood samples was analyzed using a SPADE-t-SNE-ACCENSE pipeline. Due to t-SNE limitations the dataset had to be downsampled by 57.7% (Fig. 2a) where it was decided to equal the number of cells from blood and intestinal samples for a balanced comparison, which led to the exclusion of more cells from the blood samples. Moreover, ACCENSE clustered only 50% of the t-SNE-embedded data into subsets (Fig. 2a). Together this excluded 78.8% of the cells from the analysis. The remaining 1.1 million cells were annotated into 142 phenotypically distinct immune subsets^14^ (Fig. 2a).

**Figure 2.**
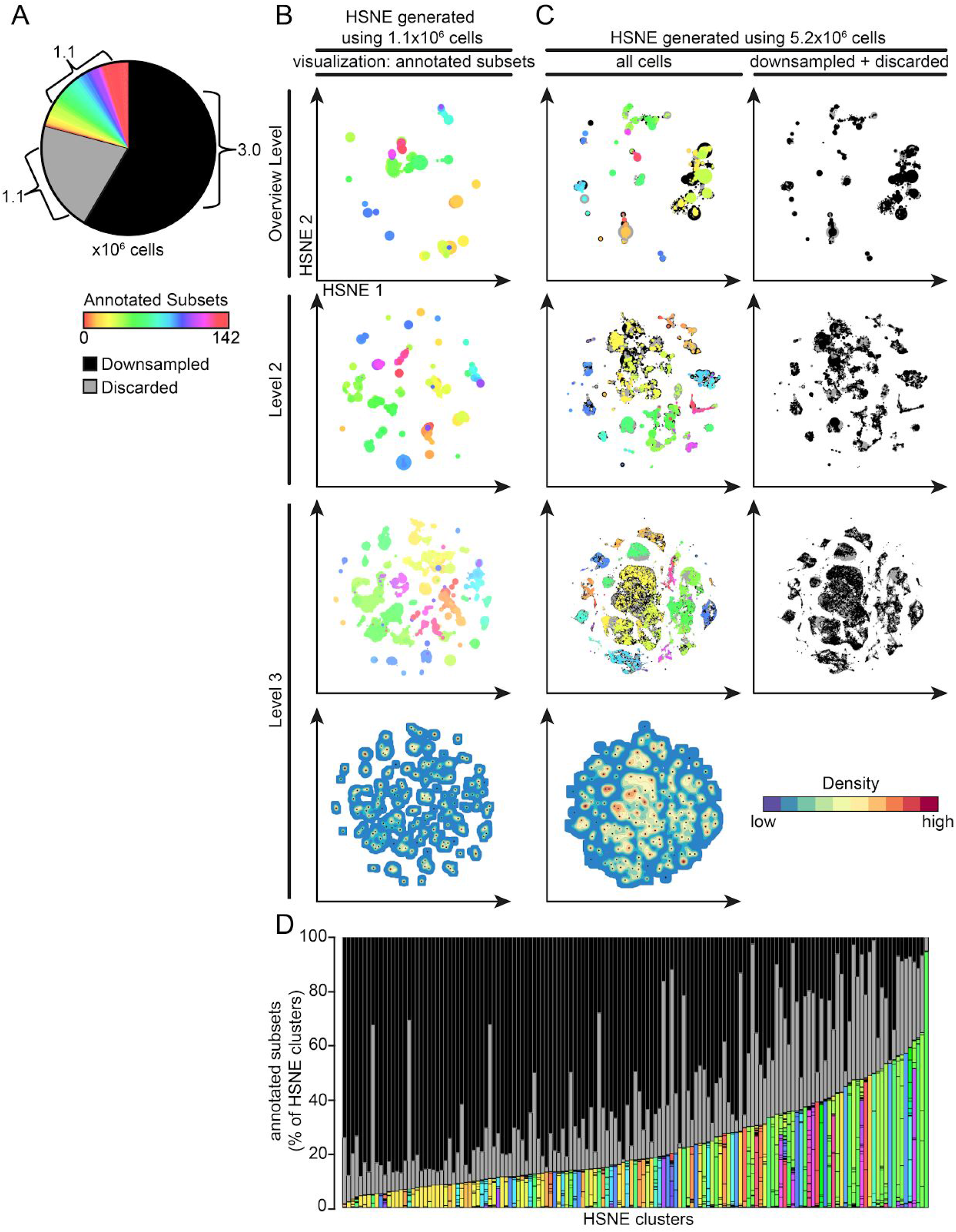
Gain of information by analyzing mass cytometry data at full resolution with Cytosplore^+HSNE^. **(a)** Pie chart showing cellular composition of the mass cytometry dataset. Color represents the subsets (N=142), as identified in our previous study^14^. Black represents the cells discarded by stochastic downsampling and grey the cells discarded by ACCENSE clustering. **(b)** Embeddings of the 1.1 million cells annotated in ref^14^ showing the top three levels of the HSNE hierarchy (five levels in total). Color represents annotations as in panel a. Size of the landmarks is proportional to the number of cells in the AoI that each landmark represents. Bottom map shows density features depicting the local probability density of cells for the level 3 embedding, where black dots indicate the centroids of identified cluster partitions using GMS clustering. **(c)** Embeddings of all 5.2 million cells, again showing only the top three levels of the hierarchy (five levels in total). Colors as in panel a. Right panels visualize landmarks representing cells discarded by stochastic downsampling (black) and the cells discarded by ACCENSE (grey). Bottom map shows density features for the level 3 embedding as described in panel b. **(d)** Frequency of annotated cells for 145 clusters identified by Cytosplore^+HSNE^ at the third hierarchical level using GMS clustering in panel c. Color coding as in panel a.

To determine whether Cytosplore^+HSNE^ could identify similar subsets, we embedded the 1.1 million annotated cells (Fig. 2b). Computation time was in the order of minutes and the analysis was finished within an hour, compared to eight weeks of computation in the original study. Color coding shows the grouping of subsets at all hierarchical levels. GMS clustering at the third level embedding (Fig. 2b, bottom panel) reveals that 75.5% of cells were assigned to a single subset by both methods (Supplementary Fig. 4). Hence, to reach similar results it was not necessary to explore the data at lower (more detailed) levels.

Next, we utilized Cytosplore^+HSNE^ to analyze the complete dataset on 5.2 million cells, thus including the cells that were discarded in the SPADE-t-SNE-ACCENSE pipeline. The embeddings show by color coding that subsets of the same immune lineage clustered at all three levels (Fig. 2c). More interestingly, the cells removed during downsampling (shown in black) and cells ignored during the ACCENSE clustering (shown in grey) were positioned throughout the entire map (Fig. 2c). We selected 145 clusters using GMS clustering at the third level and observed that the identified clusters contained variable numbers of downsampled and non-classified cells (Fig. 2d). These findings indicate that both the non-uniform downsampling and the cell losses during the ACCENSE clustering introduce a potential bias in observed heterogeneity in the immune system. Cytosplore^+HSNE^ overcomes this problem as it analyzes *all* cells and does so efficiently.

### Cytosplore^+HSNE^ Reveals Additional Complexity and Identifies Rare Subsets in the ILC Compartment

We illustrate an exploration workflow with Cytosplore^+HSNE^ using the dataset of 5.2 million cells^14^ (Fig. 3). At the overview level, 4,090 landmarks depict the general composition of the immune system (Fig. 3a) and color coding is applied to reveal CD-marker expression patterns on the basis of which the major immune lineages are identified (Fig. 3b). Next the CD7+CD3^-^ cell clusters were selected as indicated and a new higher resolution embedding was generated at level 3 of the hierarchy (Fig. 3c). Here, coloring of the landmarks based on marker expression (Fig. 3c, top panels) and a density plot of the embedding is shown (Fig. 3d) alongside the clinical features of the subjects from which the samples were obtained and the tissue-origin of the landmarks (Fig. 3c, bottom panels). This reveals a cluster of cells abundantly present in the intestine of patients with refractory celiac disease (RCDII). In addition, a large cluster of CD45RA+CD56+ NK cells and three distinct innate lymphoid cell (ILC) clusters with a characteristic lineage^-CD7+^CD161^+^CD127^+^ marker expression profile^19,20^ are visualized. Strikingly, a distinct population of CD7^+^CD127^-^CD45RA^-^ and partly CD56^+^ cells is found in between the NK, RCDII and ILC cell clusters.

**Figure 3.**
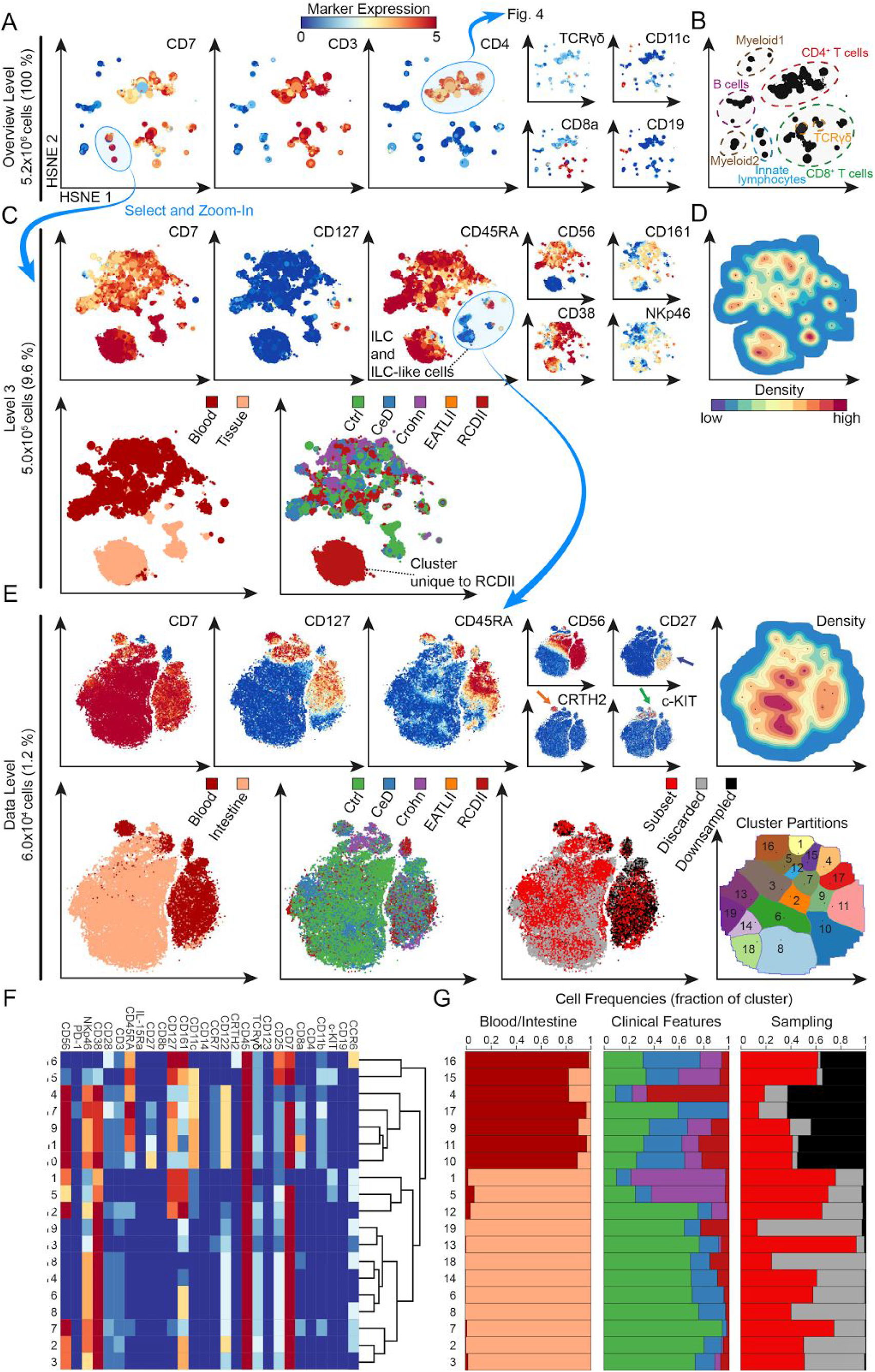
Analysis of the CD7+CD3^-^ innate lymphocyte compartment in inflammatory intestinal diseases. **(a)** First HSNE level embedding of 5.2 million cells. Color represents arcsin5-transformed marker expression as indicated. Size of the landmarks represents AoI. Blue encirclement indicates selection of landmarks representing CD7+CD3^-^ innate lymphocytes and CD4+ T cells further discussed in Figure 4. **(b)** The major immune lineages, annotated on the basis of lineage marker expression. **(c)** Third HSNE level embedding of the CD7+CD3^-^ innate lymphocytes (5.0x10^5^ cells). Color represents arcsin5-transformed marker expression in top panels, and tissue-origin and clinical features in bottom panels. Blue encirclement indicates selection of landmarks representing CD127+ILC and ILC-like cells. **(d)** Third HSNE level embedding shows density features depicting the local probability density of cells, where black dots indicate the centroids of identified cluster partitions using GMS clustering. **(e)** Embedding of the CD127+ILC and ILC-like cells (6.0x10^4^ cells) at single-cell resolution. Arrows indicate ILC1 (blue), ILC2 (orange) and ILC3 (green). Bottom-right panel shows corresponding cluster partitions using GMS clustering based on density features (top-right panel). **(f)** A heatmap summary of median expression values (same color coding as for the embeddings) of cell markers expressed by CD127+ILC and ILC-like clusters identified in panel b and hierarchical clustering thereof. **(g)** Composition of cells for each cluster is represented graphically by a horizontal bar in which segment lengths represent the proportion of cells with: (left) tissue-of-origin, (middle) disease status and (right) sampling status.

To uncover the phenotypes of these ILC-related clusters, we next embedded the ILC and ILC-like clusters (Fig. 3c, selection) at the full single-cell data level (59,775 cells; 1.2% of total) (Fig. 3e). The marker expression overlays revealed that the majority of cells are CD7^+^ and displayed variable expression levels for CD127, CD45RA and CD56 (Fig. 3e). In addition, and in line with previous reports^21,22^, (co-)expression of CD127 with CD27, CRTH2 and c-KIT revealed the phenotypes corresponding to helper-like ILC type 1, 2 and 3, respectively (indicated by arrows in Fig. 3e). Moreover, by visualizing the tissue-origin in the Cytosplore^+HSNE^ embedding the tissue-specific location of ILC and ILC-related phenotypes became evident (Fig. 3e).

Next, we performed GMS clustering on the full data level embedding which resulted in 19 phenotypically distinct clusters (Fig. 3e, right plots) based on marker expression profiles (Fig. 3f). The cell surface phenotypes of 8 out of the 19 clusters (Fig. 3f) matched previously described^21^ biological annotations (Fig. 5, black annotations) including the CRTH2+ILC2 (cluster 16), c-KIT+ILC3 (cluster 5) and CD56^-^CD127^-^lineage^-^IELs (cluster 19, 13, 18, 14, 6 and 8), the latter representing innate type of lymphocytes with dual T cell precursor and NK/ILC traits^23–25^. Remarkably, the remaining 11 clusters strongly resembled distinct ILC types, but did not fulfil the complete phenotypic requirements according to established nomenclature^21^ (Fig. 5, red annotations). For example, cluster 15 is highly similar to ILC2 (cluster 16) based on the expression of CD7, CD127, CD161 and CD25, but lacks the ILC2-defining marker CRTH2. Also, clusters 17, 9 and 11 bear close resemblance to ILC1 based on CD7+CD127+c-KIT^-^ marker expression profile, but lack the ILC-defining CD161 marker. Finally, cluster 1 is very similar to ILC3 (cluster 5) based on CD127, CD161 and c-KIT positivity, but lacks the lymphoid marker CD7. Interestingly, the ILC3 (cluster 5) and ILC3-like (cluster 1) populations resided mainly in intestinal biopsies of patient with Crohn’s disease (Fig. 3f) and may be related. Cluster 4 was mainly present in peripheral blood of patients with RCDII, suggesting a possible association with this pre-malignant disease state. Importantly, three clusters (4, 17 and 19) (Fig. 3f) were essentially missed in our previous study^14^ due to the downsampling. Finally, all identified cell clusters consist to a variable extent of cells that were downsampled in the original analysis (Fig. 3g). Thus, the analysis of the full dataset provides increased detail and confidence in establishing the phenotypes of these low abundance innate cell subsets.

### Identification of Rare CD4^+^ T Cell Subsets in Peripheral Blood

Next, we selected the CD4+ T cell lineage (Fig. 3a) and show the distribution of the landmarks at the third level, revealing several clusters within the CD4+ T cell compartment (Fig. 4a), including a small CD28^-^CD4+ T cell memory population (25,398 cells; 0.5% of total), most likely representing terminally differentiated cells^26^. Subsequent analysis at the single-cell level (Fig. 4b) identified a CD56+ population within the CD28^-^CD4+ T cells that is enriched in blood of patients with Crohn’s disease (Fig. 4b, bottom panels, dashed black circle), as well as a CD56^-^ population of CD28^-^CD4+ T cells (Fig. 4b, bottom panels, dashed yellow circle) present in blood samples of both patients and controls. Importantly, this latter cell population was not identified in our previous publication due to the non-uniform downsampling of cells (Fig. 4b).

**Figure 4.**
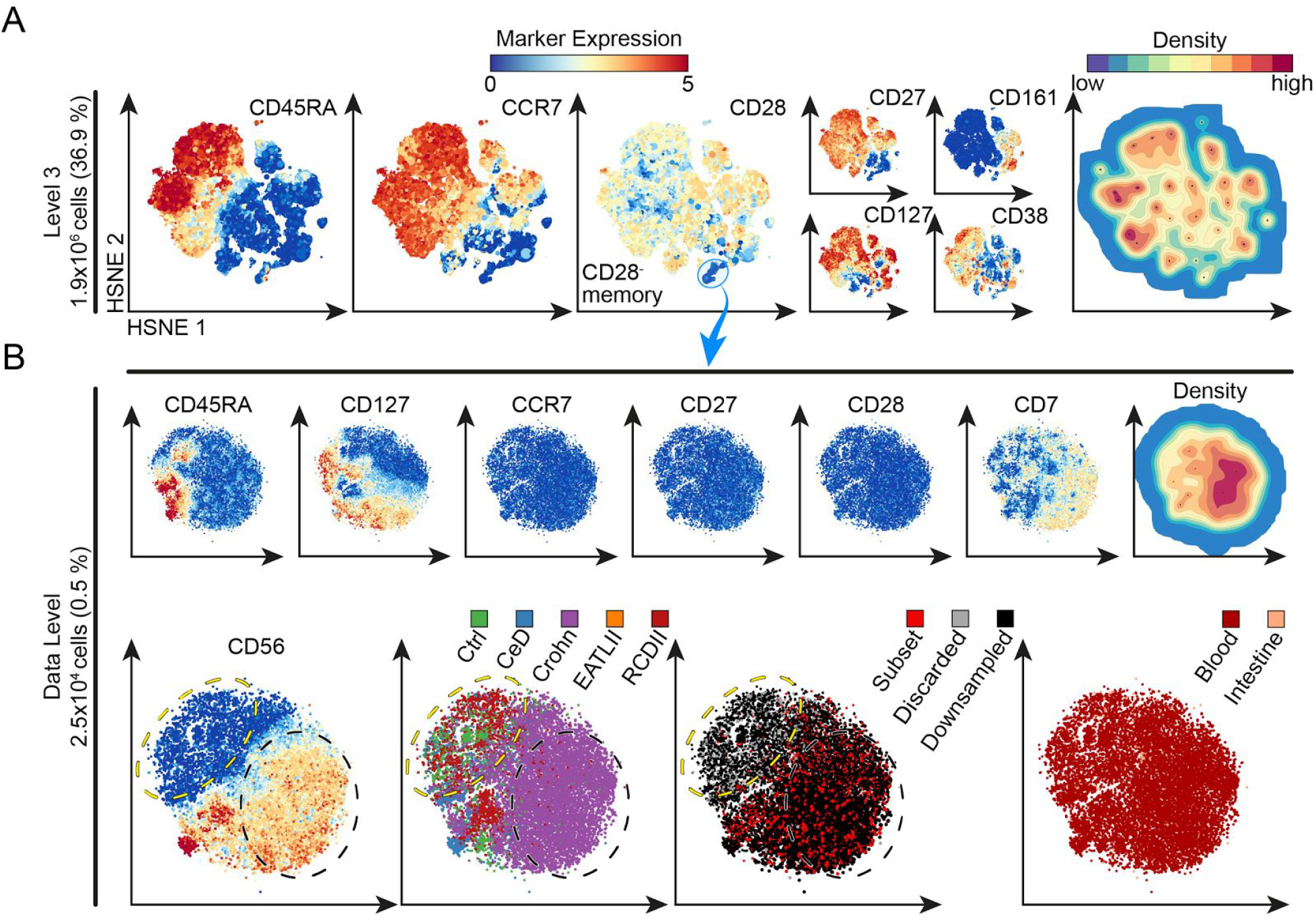
Analysis of the CD4+ T cell compartment in inflammatory intestinal diseases. **(a)** Third HSNE level embedding of the CD4+ T cells (1.4x10^6^ cells, selected in Fig. 3). Color and size of landmarks as described in Figure 3. Right panel shows density features for the level 3 embedding. Blue encirclement indicates selection of landmarks representing CD28^-^CD4+ T cells. **(b)** Embedding of the CD28^-^CD4+ T cells (2.6x10^4^ cells) at single-cell resolution. Bottom-left panel shows yellow and black dashed encirclements based on CD56^-^ and CD56+ expression, respectively. Three bottom-right panels show cells colored according to: (left) from subjects with different disease status (CeD, Crohn, EATLII, RCDII and controls), (middle) sampling status (annotated subset, discarded by ACCENSE and downsampled) and (right) tissue-of-origin (blood and intestine).

Together, these findings emphasize that Cytosplore^+HSNE^ is highly efficient in unbiased analysis of both abundant and rare cell populations in health and disease by permitting full single-cell resolution. It enables the simultaneous identification and visualization of known cell subsets and provides evidence for additional heterogeneity in the immune system, as it reveals the presence of cell clusters that were missed in a previous analysis due to downsampling of the input data. These currently unspecified cell clusters might represent intermediate stages of differentiation or novel rare cell types with presently unknown function.

### Cytosplore^+HSNE^ is Robust and Versatile Offering Advantages Over Current Single-Cell Analysis Methods

While the exploration of the hierarchy requires analysis at multiple levels, the workflow is robust and reproducible as shown in Supplementary Figure 5. In this exemplary analysis, we obtained the same Cytosplore^+HSNE^ clusters at the single-cell level upon reconstructing the hierarchy and embeddings in a matter of minutes (Methods). In addition, we tested the Cytosplore^+HSNE^ applicability to three different public mass cytometry datasets. First, we analyzed a well-characterized bone marrow dataset containing 81,747 cells^27^ as a benchmark case (Supplementary Fig. 6) and demonstrated that the landmarks in the overview level (2,632; 3.2% of total) that were selected by the HSNE algorithm were distributed across almost all of the manually gated cell types (Supplementary Fig. 6a), indicating that global data heterogeneity was accurately preserved. Also, GMS clustering resulted in HSNE clusters that were phenotypically similar to the manually gated cell types and displayed additional diversity within those subsets (Supplementary Fig. 6b). However, as the power of Cytosplore^+HSNE^ lies in its scalability to datasets exceeding millions of cells, we also tested the versatility of Cytosplore^+HSNE^ by comparing it to other state-of-the-art scalable single-cell analysis methods and accompanying large datasets (Supplementary Notes 3, Supplementary Figs. 7 and 8). Here Cytosplore^+HSNE^ computed the analyses of the VorteX dataset^5^ containing 0.8 million cells in 4 minutes, compared to 22 hours using the publicly available VorteX implementation on the same computer. Similarly, analysis of the Phenograph dataset^4^ containing 15 million cells was computed in 3.5 hours, compared to 40 hours using the publicly available Phenograph implementation on the same computer. Both analyses show that Cytosplore^+HSNE^ reproduces the main findings as presented in the original publications. More importantly, Cytosplore^+HSNE^ provides the distinct advantage of visualizing all cells and intracluster heterogeneity at subsequent levels of detail up to the single-cell level, even for the 15 million of cell dataset, without a need for downsampling. Also, VorteX failed computing the 5.2 million cell gastrointestinal dataset within 3 days of clustering (regardless of using Euclidian or Angular distance), where Cytosplore^+HSNE^ accomplished this within 29 minutes. Finally, while Phenograph did identify the rare CD56+ population of CD28^-^CD4+ T cells in the peripheral blood of individual samples (Fig. 4b), it did not reveal the association with Crohn’s disease, further highlighting the advantages of Cytosplore^+HSNE^ over these other computational tools.

**Figure 5.**
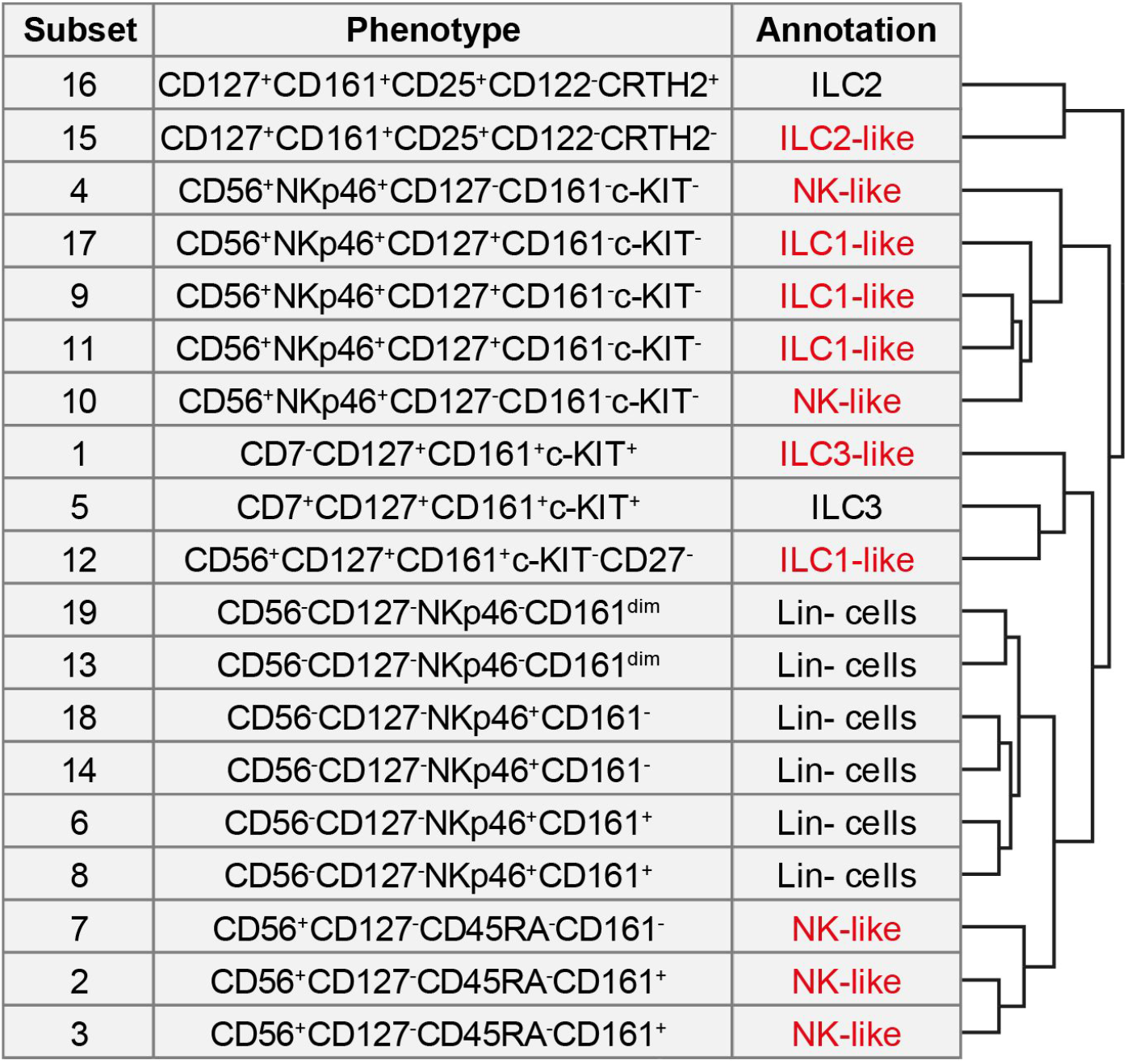
CD127+ILC and ILC-like subsets identified by Cytosplore^+HSNE^. Table showing cluster number, distinguishing phenotypic marker expression profiles and biological annotation for the clusters identified in Figure 3e. Black color indicates clusters described in previous reports and red color additional unknown clusters. Hierarchical clustering of clusters based on marker expression profile shown in the heatmap depicted in Figure 3f.

## Discussion

Mass cytometry datasets generally consist of millions of cells. Current tools can either extract global information with no single-cell resolution or provide single-cell resolution but at the expense of the number of cells that can be analyzed. Consequently, when single-cell resolution is of interest, most current tools require downsampling of datasets. However, reducing the number of included cells in the analysis pipeline may hamper the identification of rare subsets.

To overcome this problem, we introduce Cytosplore^+HSNE^. Based on a novel hierarchical embedding of the data (HSNE), Cytosplore^+HSNE^ enables the analysis of tens of millions of cells using the whole data in a fraction of the time required by currently available tools. The power of the hierarchical embedding strategy is that Cytosplore^+HSNE^ provides visualizations of the data at different levels of resolution, while preserving the non-linear phenotypic similarities of the single cells at each level. Cytosplore^+HSNE^ enables the user to interactively select groups of data points at each resolution level, either hand-picked or guided by density-based clustering, to further zoom-in on the underlying data points in the hierarchy up to the single-cell resolution. Using a dataset of 5.2 million cells we demonstrate that Cytosplore^+HSNE^ allows a rapid analysis of the composition of the cells in the dataset, that at all levels of the hierarchy the representation of these cells preserve phenotypic relationships, and that one can zoom-in on rare cell populations that were missed with other analysis tools. The identification of such rare immune subsets offers opportunities to determine cellular parameters that correlate with disease.

There is an ongoing scientific debate on the validity of clustering in t-SNE maps versus direct clustering on the high-dimensional space. However, it has been shown that stochastic neighbor embedding (SNE) preserves and separates clusters in the high dimensional space^28^. While clustering data points on highly non-linear manifolds is possible with complex models, we argue that the presented approach simplifies clustering considerably. We show that HSNE efficiently unfolds the non-linearity in the high-dimensional data, as other SNE approaches do and therefore simpler clustering methods based on locality in the map suffice to partition the data faithfully (e.g. the density-based Gaussian mean shift clustering, implemented in Cytosplore^+HSNE^). Especially when combined with an interactive quality control mechanism to visually inspect residual variance within each cluster, the kernel size can be selected such that within-cluster variance is minimized, and thereby supports the validity of the cluster with respect to potential underclustering. This is indeed confirmed by comparisons to other scalable tools (i.e. Phenograph and VorteX), showing that Cytosplore^+HSNE^ provides a superior discriminatory ability to identify and visualize rare phenotypically distinct cell clusters in large datasets in a very short time span. However, depending on user preference, Cytosplore^+HSNE^ can be used in conjunction with such direct clustering approaches. This allows the user to identify additional heterogeneity that is potentially missed by direct clustering, and provides the tools for an informed merging and splitting of clusters as the user deems appropriate. The recent application of mass cytometry and other high-dimensional single cell analysis techniques has greatly increased the number of phenotypically distinct cell clusters within the immune system. This raises obvious questions about the true distinctiveness and function of such cell clusters in health and disease, an issue that is beyond the scope of the present study but needs to be addressed in future studies.

In conclusion, Cytosplore^+HSNE^ allows an interactive and fast analysis of large high-dimensional mass cytometry datasets from a global overview to the single-cell level and is coupled to patient-specific features. This may provide crucial information for the identification of disease-associated changes in the adaptive and innate immune system which may aid in the development of disease- and patient-specific treatment protocols. Finally, Cytosplore^+HSNE^ applicability goes beyond analyzing mass cytometry datasets as it is able to analyze any high-dimensional single-cell dataset.

## Methods

### HSNE algorithm

Hierarchical Stochastic Neighbor Embedding (HSNE) builds a hierarchy of local and non-linear similarities of high-dimensional data points^13^, where landmarks on a coarser level of the hierarchy represent a set of similar points or landmarks of the preceding more detailed level. To represent the non-linear structures of the data, the similarity of these landmarks is not described by Euclidian distance, but by the concept of area of influence (AoI) on landmarks of the preceding level. The similarities described in every level of the hierarchy are then used as input for an adapted version of the similarity-based embedding technique BH-SNE^15^ for visualization. The algorithm works as follows: First a weighted k-nearest neighbor (kNN) graph is computed from the raw input data. For optimal performance and scalability the neighborhoods are approximated as described in ref.^16^. The weight of the link between two data points in the kNN graph describes the similarity of the connected data points.

In the subsequent steps the hierarchy is built, based on the similarities of the data level. To this extent, a number of random walks of predefined length is carried out starting from every node in the kNN graph, using the similarities as probability for the next jump; similar nodes to the current node are more likely to be the target of the next jump. Nodes in the graph that are reached more often are considered more important and selected as landmarks for the next coarser level. The number of landmarks is selected in a data-driven fashion, based on this importance. The AoI of a landmark is defined by a second set of random walks started from all nodes (data points or landmarks on the preceding level). Here, the length is not predefined. Rather, once a landmark is reached the random walk terminates. The influence on the node is then defined for every reached landmark as the fraction of walks that terminated in that landmark. Inversely, the AoI for each landmark is defined as the set of all nodes that reached this landmark at least once in this second set of random walks. Consequently, since multiple random walks initiated at the same node can end in different nodes, the AoIs of different landmarks can overlap. Consequently, through the similarity of two landmarks, their connection in the neighborhood graph is defined by the overlap of their corresponding AoIs, weighted by the influence defined on each node within that overlap. This process is carried out iteratively, until a predefined number of hierarchical levels has been constructed. For the full technical details we refer to our previous work^13^.

### HSNE implementation in Cytosplore^+HSNE^

We implemented our integrated analysis tool Cytosplore^+HSNE^ using a combination of C++, javascript and OpenGL. All computationally demanding parts are implemented in C++ and make use of parallelization, where possible. The density estimation and GMS clustering make use of the graphics processing unit (GPU), as described in our original publication on Cytosplore^29^, if possible, allowing clustering of millions of points in less than a second. We implemented the visualizations of the embedding in OpenGL on the GPU, for optimal performance, and less computational demanding visualizations, such as the heatmap, in javascript. We implemented the HSNE algorithm in C++, as presented in ref.^13^. Since we use sparse data structures, memory consumption strongly depends on data complexity. Maximum memory consumption during the construction of a four level hierarchy plus overview embedding of the 841,644 cell VorteX dataset was 1,684 MB, construction of a five level hierarchy of our human inflammatory intestinal diseases dataset, consisting of 5,220,347 cells required a maximum of 9,357 MB of main memory, and finally, the 15,299,616 cell Phenograph dataset required a maximum of 24.3 GB of memory during the computation of a five level hierarchy plus the overview embedding. Computation times for the described hierarchies plus the first level embedding after 1,000 iterations were 4 minutes, 29 minutes, and, 3 hours and 37 minutes, respectively, on a HP Z440 workstation with a single intel Xeon E5-1620 v3 CPU (4 cores) clocked at 3.5 Ghz, 64 GB of main memory and an nVidia Geforce GTX 980 GPU with 4 GB of memory, running Windows 7.

#### Code availability

Upon peer-reviewed journal publication of this manuscript, we provide a Cytosplore^+HSNE^ installer for Windows, allowing exploration of several million cells, for academic use at http://www.cytosplore.org.

### Human gastrointestinal disorders mass cytometry dataset

Detailed description of the mass cytometry dataset on human gastrointestinal disorders can be found in our previous work^14^. In brief, samples (N=102) were collected from patients who were undergoing routine diagnostic endoscopies. Cells from the epithelium and lamina propria were isolated from two or three intestinal biopsies by treatment with EDTA followed by a collagenase mix under rotation at 37° C. We analyzed single-cell suspensions from biological samples including duodenum biopsies (N=36), rectum biopsies (N=13), perianal fistulas (N=6), and PBMC from control individuals (N=15) and from patients with inflammatory intestinal diseases (celiac disease (CeD), N=13; refractory celiac disease type II (RCDII), N=5; enteropathy-associated T cell lymphoma type II (EATLII), N=1 and Crohn’s disease (Crohn), N=10). A CyTOF panel of 32 metal isotope-tagged monoclonal antibodies was designed to obtain a global overview of the heterogeneity of the innate and adaptive immune system. Primary antibody metal-conjugates were either purchased or conjugated in-house. Procedures for mass cytometry antibody staining and data acquisition were carried out as previously described^27^. CyTOF data were acquired and analyzed on-the-fly, using dual-count mode and noise-reduction on. All other settings were either default settings or optimized with a tuning solution. After data acquisition, the mass bead signal was used to normalize the short-term signal fluctuations with the reference EQ passport P13H2302 during the course of each experiment and the bead events were removed^30^.

Upon publication in a peer-reviewed journal, the dataset will be made publicly available on Cytobank, experiment no 60564. https://community.cytobank.org/cytobank/experiments/60564

### Processing of mass cytometry data

We transformed data from the human inflammatory intestinal diseases dataset using hyperbolic arcsin with a cofactor of 5 directly within Cytosplore^+HSNE^. We discriminated live, single CD45+ immune cells with DNA stains and event length for the human inflammatory intestinal diseases study. We analyzed other data (Phenograph and VorteX datasets) as was available, except the transformation using hyperbolic arcsin with a cofactor of 5.

### Cytosplore^+HSNE^ Analysis

Cytosplore^+HSNE^ facilitates the complete exploration pipeline in an integrated fashion (see Supplementary Video 1). All presented tools are available for every step of the exploration and every level of the hierarchy. Data analysis in Cytosplore^+HSNE^ included the following steps: We applied the arcsin transform with a cofactor of five upon loading the datasets. After that we started a new HSNE analysis and defined the markers that should be used for the similarity computation. We used markers CD3, CD4, CD7, CD8a, CD8b, CD11b, CD11c, CD14, CD19, CD25, CD27, CD28, CD34, CD38, CD45, CD45RA, CD56, CD103, CD122, CD123, CD127 CD161, CCR6, CCR7, c-KIT, CRTH2, IL-15Ra, IL-21R, NKp46, PD-1, TCRab, and TCRgd for the human inflammatory intestinal diseases dataset, all available markers for the bone marrow benchmark dataset, surface markers CD3, CD7, CD11b, CD15, CD19, CD33, CD34, CD38, CD41, CD44, CD45, CD47, CD64, CD117, CD123 and HLA-DR for the Phenograph dataset, and markers CD3, CD4, CD5, CD8, CD11b, CD11c, CD16/32, CD19, CD23, CD25, CD27, CD34, CD43, CD44, CD45.2, CD49b, CD64, CD103, CD115, CD138, CD150, 120g8, B220, CCR7, c-KIT, F4/80, FceR1a, Foxp3, IgD, IgM, Ly6C, Ly6G, MHCII, NKp46, Sca1, SiglecF, TCRb, TCRgd and Ter119 to construct the hierarchy for the VorteX dataset. We used the standard parameters for the hierarchy construction; number of random walks for landmark selection: N=100, random walk length: L=15, number of random walks for influence computation: N=15. For any clustering that occurred the GMS grid size was set to S=256^2^. The reduction factor from one level in the hierarchy to the next coarser level is completely data-driven. In our experiments with mass cytometry data the number of landmarks was consistently reduced by roughly one order of magnitude from one level to the next. Embeddings consisting of only a few hundred points usually provide little insight. Therefore we defined the number of levels such that the overview level could be expected to consist of in the order of 1,0 landmarks meaning N=5 for the human inflammatory intestinal diseases dataset and Phenograph dataset, N=3 for the bone marrow benchmark dataset, and N=4 for the VorteX dataset. Building the hierarchy automatically creates a visualization of the overview level using BH-SNE. Cytosplore^+HSNE^ enables color coding of the landmarks using expression (e.g. Fig. 3a) of any provided markers or by sample. For example, we created the clinical feature (e.g. Fig. 3c, bottom-left panel) and blood/intestine (e.g. Fig. 3c, bottom-right panel) color schemes based on samples for the human inflammatory intestinal diseases dataset within Cytosplore^+HSNE^, and for the Phenograph dataset we created a color scheme that represented the sample coloring as provided in ref.^4^ (Supplementary Fig. 7). For zooming into the data we generally selected cells based on visible clusters, either using manual selection or by selecting clusters derived by using the GMS clustering. For the VorteX dataset we clustered the third level embedding (Supplementary Fig. 8). We specified a kernel size of 0.18 of the embedding size, to match the 48 clusters created by the X-shift clustering described in ref.^5^, resulting in 50 clusters.

For subset classification we first cluster the embedding at a given level using the GMS clustering. Next, we inspect the clustering by using the integrated descriptive marker statistics and heatmap visualization. If there is still meaningful variation of the marker expression within clusters we zoom further into these clusters. If clusters are phenotypically homogeneous the corresponding cell types are defined by inspecting the full marker expression profile in the heatmap and then the cluster is exported from any level in the hierarchy.

## Acknowledgements

The research leading to these results has received funding from Leiden University Medical Center, the Netherlands Organization for Scientific Research (ZonMW grant 91112008) and the Technology Foundation STW, the Netherlands (VAnPIRe; grant 12720, and Genes in Space; grant 12721).

We thank Drs. M.W. Schilham, M. Yazdanbakhsh, J Goeman, K Schepers, J van Bergen and S.E. de Jong for critical review of the manuscript and B. van Lew for narrating the Supplemental Video.

## Author Contributions

V.v.U., T.H., N.P., F.K., A.V. and B.L. conceived the study. T.H., N.P., A.V., and B.L. developed the HSNE method and implementation in Cytosplore^+HSNE^ V.v.U. and F.K. performed biological analysis and interpretation. T.H. performed the t-SNE scalability analysis and comparison. V.v.U. performed the hierarchy robustness analysis. V.v.U. and T.H. performed the comparison with other methods. N.L., M.R. and E.E. provided conceptual input. V.v.U., T.H., N.P., F.K., A.V. and B.L. wrote the manuscript. All authors discussed the results and commented on the manuscript.

## Supplementary Material

### Additional Supplementary Notes

#### 1. Cytosplore^+HSNE^ is reproducible and robust

Cytosplore^+HSNE^ allows significant user interaction during the exploration of the HSNE hierarchy, where the embedding visualizations and integrated clustering provide strong guidance. Independent explorations of the 5.2 million dataset, following the same zooming-in strategy are shown in Supplementary Figure 5. While the embeddings slightly vary at all levels, (mostly in rotation and reflection of the map), the same high level structure is found in all explorations. The robust separation of these structures guides the user in the selection and zooming-in process, resulting in highly similar embeddings down to the data level.

Focusing on separate regions of the data and interactively zooming into these separately provides significantly more detail than is possible by direct dimensionality reduction or clustering of the complete dataset (Figs. 3 and 4). However, Cytosplore^+HSNE^ does provide the possibility to visualize the complete dataset at the data level (Supplementary Fig. 1a). A dataset consisting of 1 million cells created by randomly sampling the 5.2 million cell dataset presented in the main text and three smaller ones derived from this were analysed with HSNE and t-SNE resulting in highly similar embeddings (Supplementary Fig. 1a).

Supplementary Figure 1b shows the robustness of HSNE with regard to downsampling as well as the superiority of the HSNE data reduction towards the overview level, compared to random downsampling. Here the embeddings within each column are similar, indicating that HSNE captures similar features even with downsampled data. However, detail increases with growing data sizes even if the number of landmarks are comparable between datasets. Thus the HSNE hierarchy preserves the non-linear structures in the data when reducing the data for visualization at the more abstract levels, while these structures can be lost during random downsampling.

The difference in detail is especially striking when comparing the complete HSNE hierarchy of 1 million cells (Supplementary Fig. 1b, top row) to the t-SNE embeddings of randomly sampled datasets of similar sizes as the HSNE levels (Supplementary Fig. 1a, bottom row).

#### 2. Millions of cells cause performance issues and overcrowding in t-SNE

Although feasible with a strong computational infrastructure, t-SNE suffers from several problems when analyzing datasets exceeding hundreds of thousands of cells. Three main parameters influence the result of a t-SNE embedding: the number of iterations for the gradient descent *i*, perplexity *p* and theta *t* (the latter only for BH-SNE). Cytobank provides a brief analysis of the parameters^1^ that shows diminishing returns for p and t, beyond certain values, which can sensibly be used as defaults and do not significantly change with the input data size. In contrast, i needs to be adjusted with increasing data sizes. We show that the commonly used default value of i=1,000 is not enough to properly embed millions of cells (Supplementary Fig. 2). All embeddings were created using A-tSNE, implemented in Cytosplore, using the default parameters of p=30 and t=0.01. Supplementary Figure 2a-c show embeddings of 1 million, 2 million and 5 million cells, respectively, randomly sampled from the 5.2 million cell dataset presented in the main text after 1,000 iterations. Computation time for the embeddings were (a) 5.5 h, (b) 13 h, and (c) 54 h. Supplementary Figure 2d-f show the same embeddings after 4,000 further iterations. Total computation time for the embeddings were (d) 19.5 h, (e) 45.5 h, and (f) 252 h.

While Supplementary Figure 2a seems to provide a good separation for some high level clusters Supplementary Figure 2b and c show typical artifacts of a non-converged embedding, i.e. the cells concentrate strongly in the center of the visualization, often forming a cross shape along the two axes as is clearly visible in the density plots.

All embeddings evolved significantly after 4,000 additional iterations (Supplementary Fig. 2d-f), indicating that 1,000 iterations are not enough to fully converge for these large data sizes. Even after 5,000 iterations and 252 h of computation Supplementary Figure 2f still shows similar artifacts. Another problem of computing t-SNE for such large datasets is overcrowding. All embeddings show signs of overcrowding. Only large scale neighborhoods can be identified in Supplementary Figure 2d, while structure within these neighborhoods is hard to identify due to the large number of cells, even in the density plot. Also, in Supplementary Figure 2e and **f** some ‘color smear’ is present in the single-cell plots indicating that local neighborhoods were not resolved properly by the t-SNE algorithm. Intuitively, t-SNE accounts for small neighborhoods. By increasing the size of the input data local neighborhoods will often become less strongly connected and can tear, resulting in the displacement of cells in the plot. These effects might be reduced by increasing the perplexity value^2^.

Increasing p will help in the separation of high level clusters, however, at the cost of intracluster separation, as there will be less visual space for each cluster. A detailed analysis of the neighborhood conservation of different dimensionality reduction techniques, including t-SNE, can be found in our previous work^13^.

#### 3. Cytosplore^+HSNE^ offers advantages over current scalable single-cell analysis methods

We investigated the generalizability as well the scalability of Cytosplore^+HSNE^ by comparison to two other state-of-the-art scalable single-cell analysis methods and accompanying public datasets (Phenograph and VorteX). Both techniques use a clustering method followed by visualization of the generated clusters.

Phenograph achieves this by the Louvain community detection method for partitioning of the kNN graph, followed by a t-SNE embedding of the communities based on their median values. The resulting embedding places the communities in a global context, but cannot display the details of the single-cell complexity within the communities. Using Cytosplore^+HSNE^ we were able to reproduce the clusters of the Phenograph bone marrow dataset, consisting of 15 million cells, after 3.5 hours of computation, compared to 40 hours with the Phenograph algorithm (clustering per individual samples) on the same computer. Also, Cytosplore^+HSNE^ only required 29 minutes to compute the 5.2 million cell gastrointestinal dataset, while Phenograph required 4 hours. In addition to the significantly faster computation, Cytosplore^+HSNE^ provides the distinct advantage of visualizing all cells and intracluster heterogeneity at subsequent levels of detail (Supplementary Fig. 6).

VorteX first clusters the data using the X-shift algorithm, and then visualizes the result by random sampling of cells from the clusters for visualization in a single-cell force-directed layout. The sampling is necessary, as the force-directed layout can computationally handle 30,000 cells only. Therefore, the resulting single-cell visualization shows only 3.6 % of the original dataset. Although the technique allows for more detailed cellular visualization compared to Phenograph, a time-consuming second computation is required for every additional analysis on individual immune lineages. In a direct comparison Cytosplore^+HSNE^ recapitulated the murine bone marrow clusters at the second level of a 4 level hierarchy in 4 minutes while VorteX required 22 hours (Supplementary Fig. 7a, b). In addition, by applying the zooming-in approach, we obtained the single-cell details for the plasmacytoid dendritic cell lineage within seconds (Supplementary Fig. 7c). Finally, VorteX failed computing the 5.2 million cell gastrointestinal dataset within 3 days of clustering (regardless of using Euclidian or Angular distance).

**Supplementary Figure 1.**
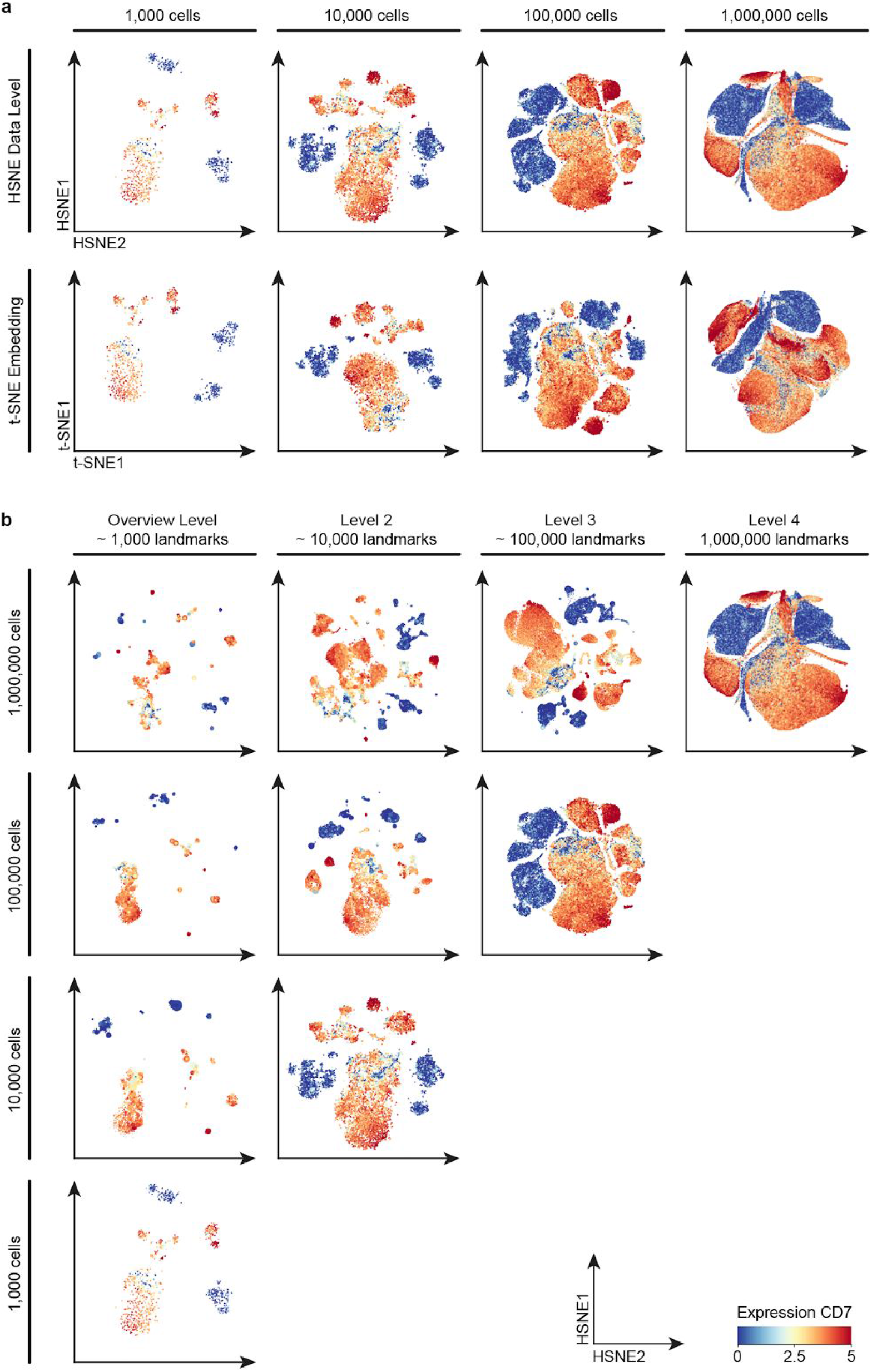
Comparison of robustness with regard to downsampling between t-SNE and HSNE. (**a**) Comparison of t-SNE (bottom row) and HSNE (top row) data level embeddings for datasets of different sizes (columns). First, 1 million cells were randomly sampled from the 5.2 million cell dataset, the smaller datasets were then created by randomly sampling the next largest one. All plots were created after 1,000 iterations. The 1 million cell embeddings were not fully converged. Color indicates CD7 expression. (**b**) Robustness of the HSNE hierarchy with regard to downsampling. Each row shows the datasets as described above. Embeddings for the complete hierarchy of log10(N / 100) levels, with N being the number of cells, are shown in the columns. Color as in panel a. Numbers of landmarks are approximated, indicating a reduction of one order of magnitude per level. In all columns the amount of detail increases towards the top (larger datasets), even though all embeddings in a column consist of roughly the same number of points. This implies that the preservation of non-linear neighborhoods by HSNE conserves structure that is lost by random downsampling.

**Supplementary Figure 2.**
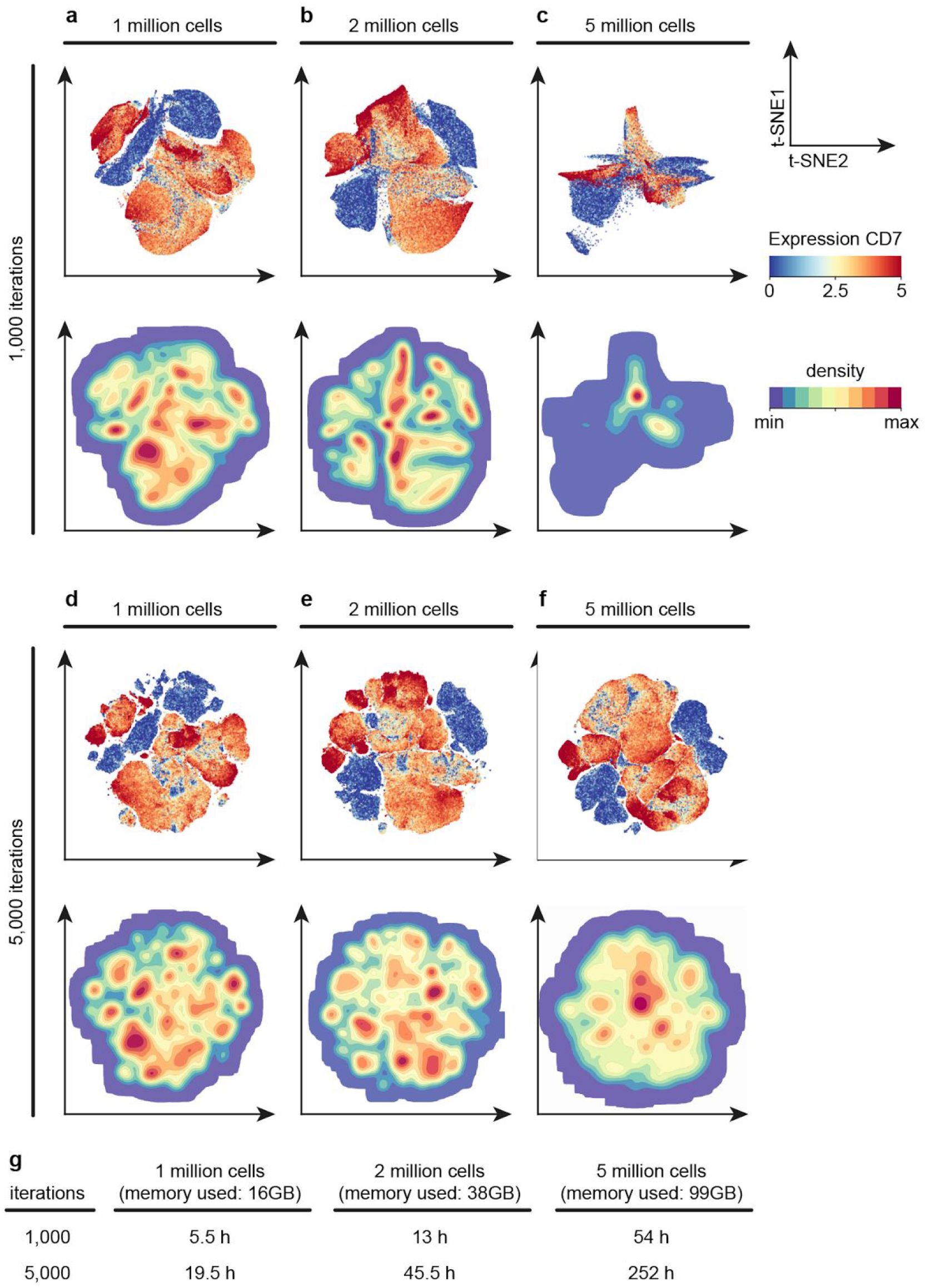
t-SNE embeddings of millions of cells show overcrowding and artifacts caused by insufficient optimization. (**a-c**) Single-cell (top row) and density-based (bottom row) visualizations of t-SNE embeddings of (**a**) 1, (**b**) 2 and (**c**) 5 million cells, respectively, after 1,000 iterations, the standard setting used in many t-SNE applications. Color in the single-cell visualization corresponds to the CD7 marker expression; in the density visualization to the cell density in the t-SNE plot. (**d-f**) The same embeddings, consisting of (**d**) 1, (**e**) 2 and (**f**) 5 million cells, respectively, after 4,000 additional iterations, resulting in a total of 5,000 iterations. Colors as above. (**g**) Computation times for the different t-SNE computations.

**Supplementary Figure 3.**
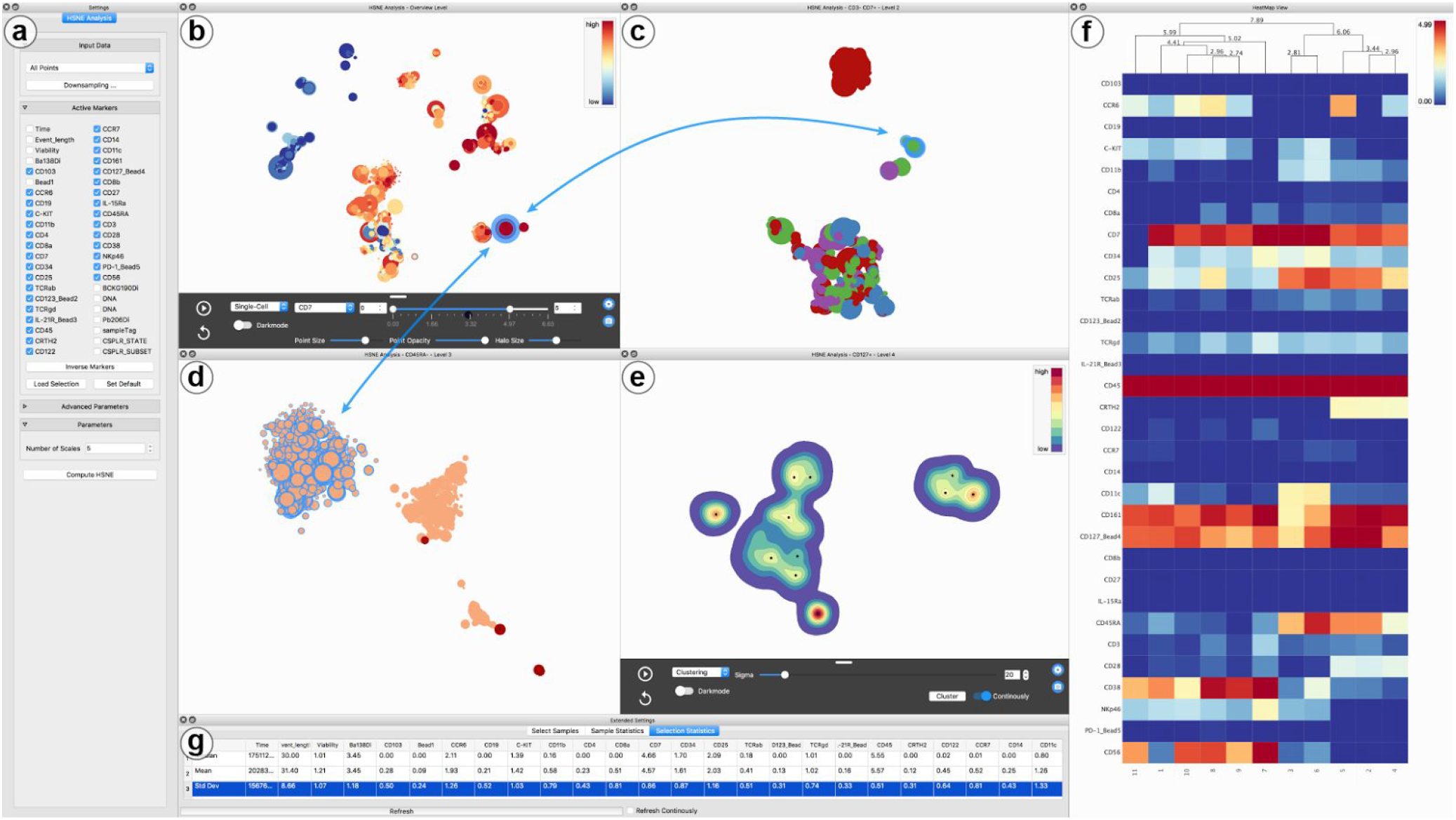
The Cytosplore^+HSNE^ software. (**a**) Settings panel for the HSNE analysis. (**b-e**) Zoom into the Innate Lymphocytes as shown in Figure 2 and Supplementary Figure 3. (**b**) overview level, (**c**) level 2, (**d**) level 3, (**e**) level 4. Color shows; (**b**) CD7 marker expression, (**c**) clinical features, (**d**) tissue origin, (**e**) cell density. A selection in panel d is highlighted in panel b,c, and d by blue halos around circles and arrows. Note, arrows added for clarity only and are not part of the software. (**f**) heatmap visualization of the median values of the clusters generated by GMS clustering based on the density visualization in panel e. Color shows marker expression. (**g**) Statistics of the selection shown in panel b-d.

**Supplementary Figure 4.**
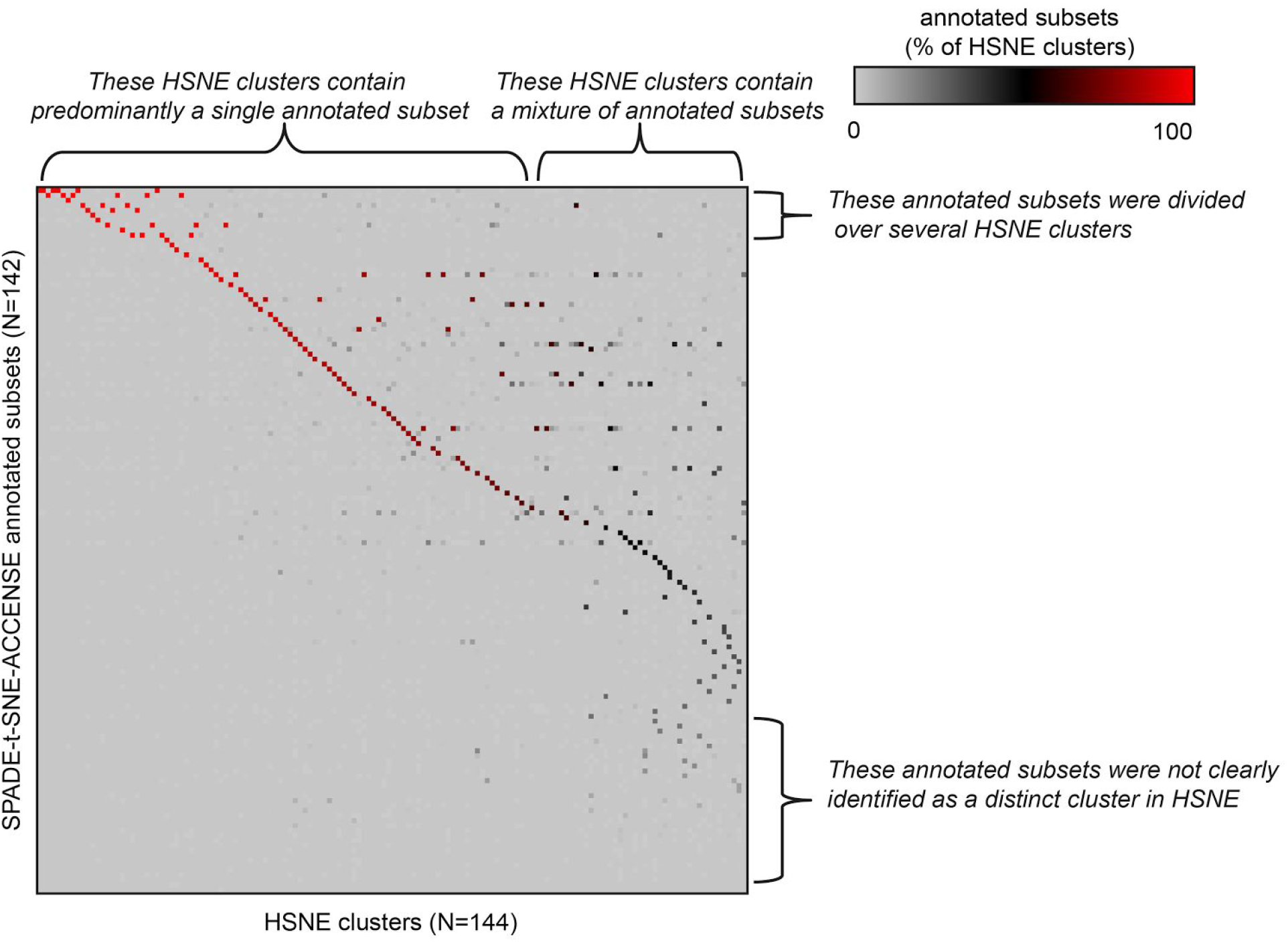
Comparisons of cellular composition of the clusters identified with Cytosplore^+HSNE^ with the previously annotated subsets using the SPADE-t-SNE-ACCENSE method. Rows indicate the individual SPADE-t-SNE-ACCENSE annotated subsets (N = 142) identified in the previous study^14^ (N = 142) and columns indicate the individual clusters identified with Cytosplore^+HSNE^ (N = 144) of the same 1.1 million cells from the gastrointestinal dataset. Color indicates the fraction of the cluster containing cells assigned to a single subset as annotated with SPADE-t-SNE-ACCENSE.

**Supplementary Figure 5.**
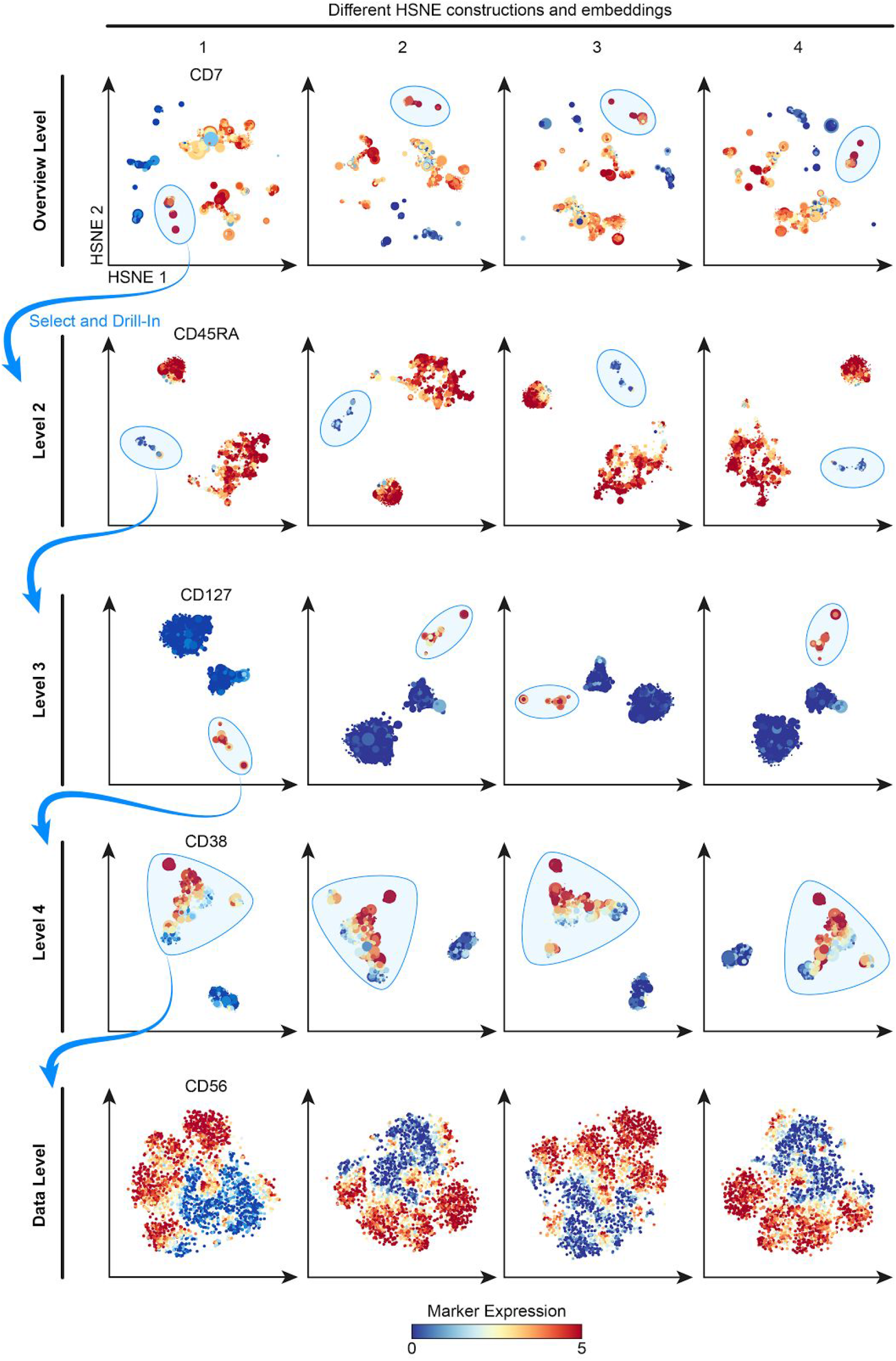
Reproducibility of the hierarchy and the embeddings. Four independent Cytosplore^+HSNE^ analyses are shown (columns) reproducing the hierarchy construction and exploration of the data with the same zooming-in strategy (blue encirclements). Color-coding indicates arcsin5-transformed marker expression.

**Supplementary Figure 6.**
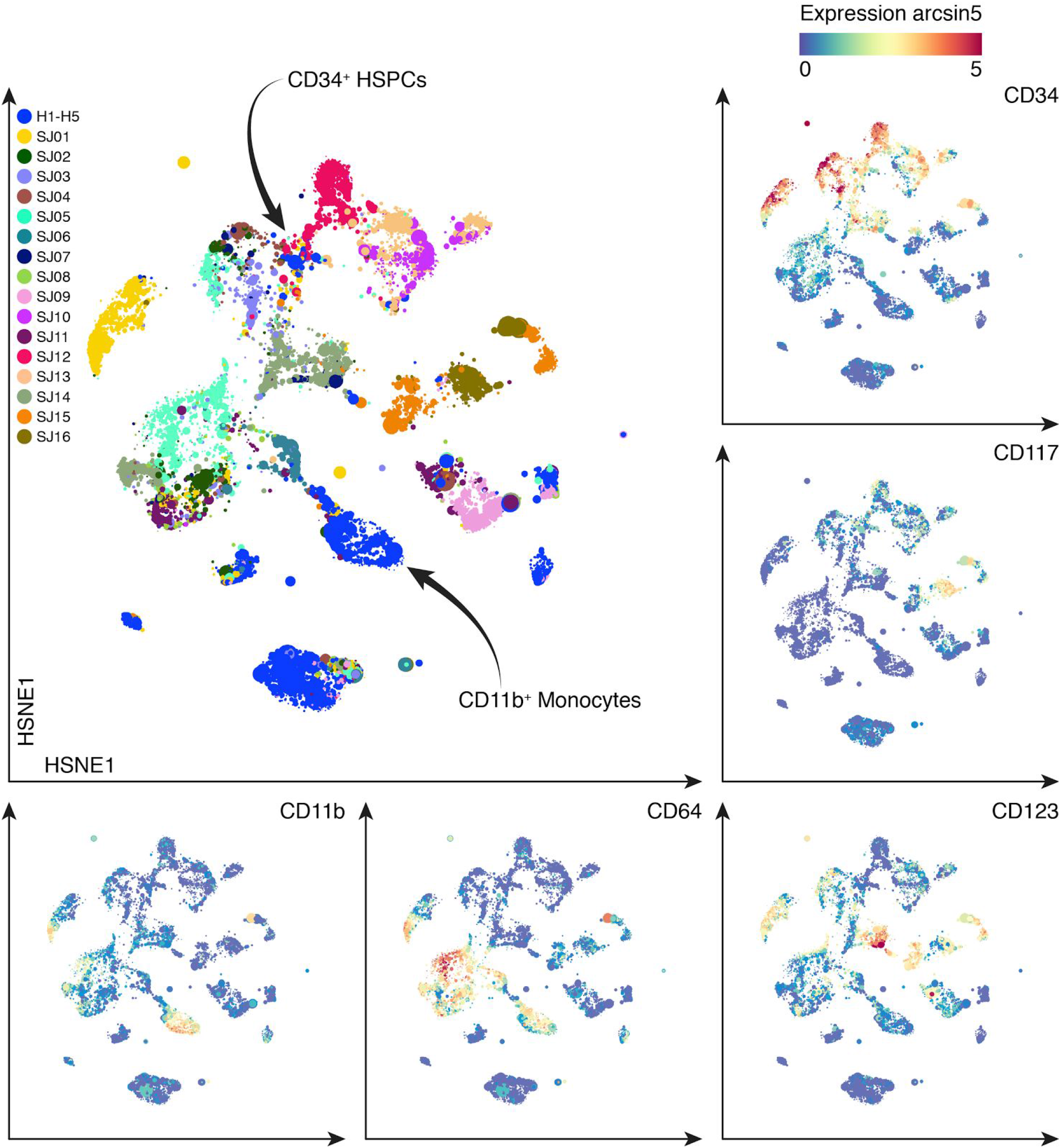
Cytosplore^+HSNE^ analysis of the Phenograph bone marrow dataset. Cytosplore^+HSNE^ embeddings of the full 15.0 million cells of the Phenograph human bone marrow dataset (overview level of a 5 level hierarchy). Color coding of main panel (top left) by patient identity. In additional panels, color coding indicates arcsin5-transformed marker expression. The above shows a comparison with Figure 3 of the original study^4^.

**Supplementary Figure 7.**
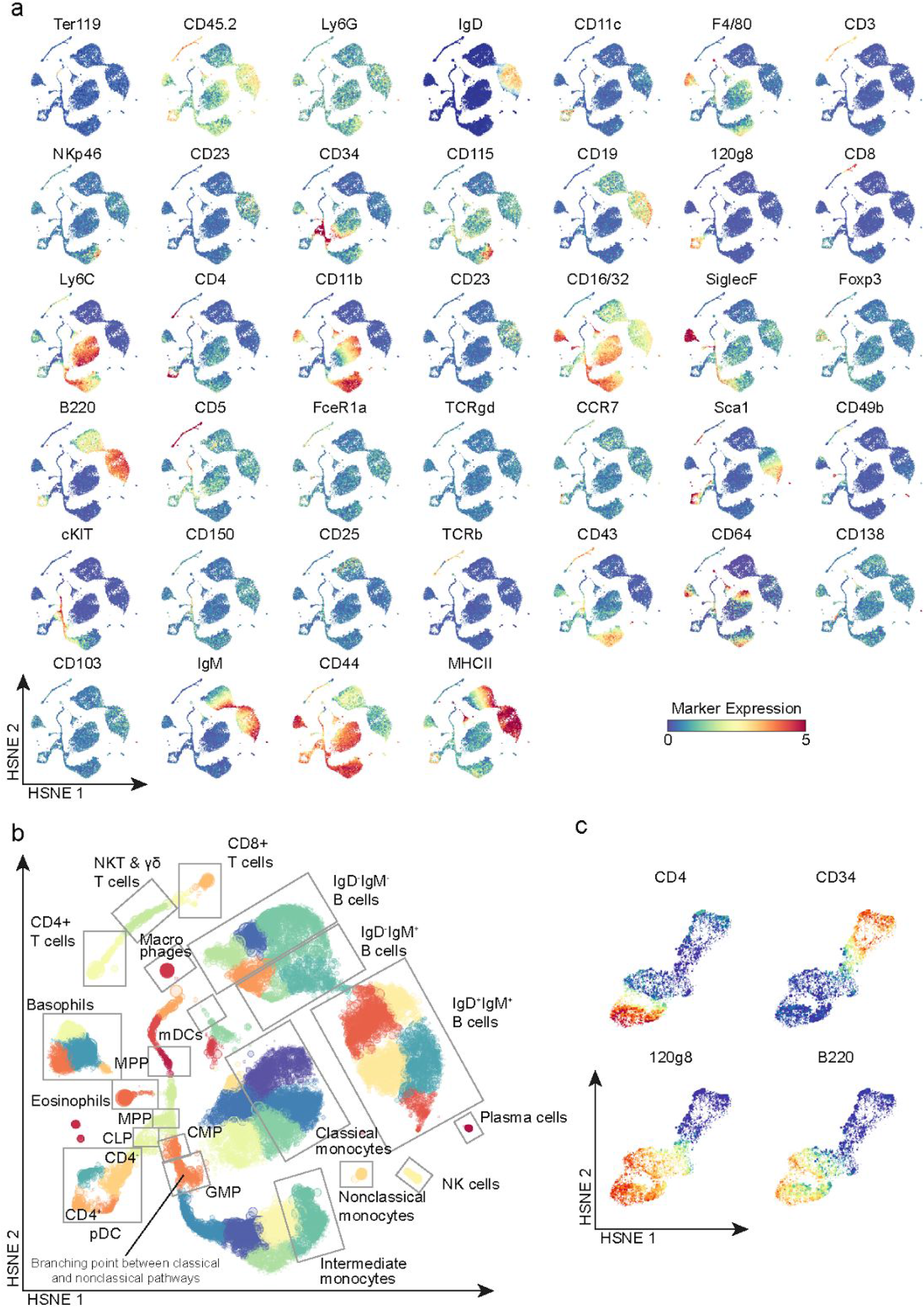
Cytosplore^+HSNE^ analysis of the VorteX bone marrow dataset. (**a**) Cytosplore^+HSNE^ embeddings of the full 0.8 million cells of the VorteX mouse bone marrow dataset (2^nd^ hierarchical level of 4 in total). Color coding indicates arcsin5-transformed marker expression. (**b**) Embedding as in panel a. Color coded for 50 clusters identified with Cytosplore^+HSNE^ Shaded boxes show locations of hand-gated cell populations. (**c**) Embeddings of zoomed-in populations related to pDC development (3^rd^ hierarchical level of 4 in total). The above shows a comparison with Figure 2 of the original study^5^.

## Footnotes

1 https://support.cytobank.org/hc/en-us/articles/206439707-How-to-Configure-and-Run-a-viSNE-Analysis#iterations

2 http://blog.cytobank.org/2017/01/17/fine-tune-visne-to-get-the-most-of-your-single-cell-data-analysis/

## References

1. Saeys, Y., Gassen, S. V. & Lambrecht, B. N. Computational flow cytometry: helping to make sense of high-dimensional immunology data. Nat. Rev. Immunol. 16, 449–462 (2016).

2. Qiu, P. et al. Extracting a cellular hierarchy from high-dimensional cytometry data with SPADE. Nat. Biotechnol. 29, 886–891 (2011).

3. Zunder, E. R., Lujan, E., Goltsev, Y., Wernig, M. & Nolan, G. P. A continuous molecular roadmap to iPSC reprogramming through progression analysis of single-cell mass cytometry. Cell Stem Cell 16, 323–337 (2015).

4. Levine, J. H. et al. Data-Driven Phenotypic Dissection of AML Reveals Progenitor-like Cells that Correlate with Prognosis. Cell 162, 184–197 (2015).

5. Samusik, N., Good, Z., Spitzer, M. H., Davis, K. L. & Nolan, G. P. Automated mapping of phenotype space with single-cell data. Nat. Methods 13, 493–496 (2016).

6. Spitzer, M. H. et al. IMMUNOLOGY. An interactive reference framework for modeling a dynamic immune system. Science 349, 1259425 (2015).

7. Hotelling, H. Analysis of a Complex of Statistical Variables Into Principal Components. (1933).

8. van der Maaten LJP and Hinton, G. E. Visualizing High-Dimensional Data Using t-SNE. J. Mach. Learn. Res. 2579–2605 (2008).

9. Amir, E.-A. D. et al. viSNE enables visualization of high dimensional single-cell data and reveals phenotypic heterogeneity of leukemia. Nat. Biotechnol. 31, 545–552 (2013).

10. Haghverdi, L., Buettner, F. & Theis, F. J. Diffusion maps for high-dimensional single-cell analysis of differentiation data. Bioinformatics 31, 2989–2998 (2015).

11. Bendall, S. C., Nolan, G. P., Roederer, M. & Chattopadhyay, P. K. A deep profiler’s guide to cytometry. Trends Immunol. 33, 323–332 (2012).

12. Chattopadhyay, P. K., Gierahn, T. M., Roederer, M. & Love, J. C. Single-cell technologies for monitoring immune systems. Nat. Immunol. 15, 128–135 (2014).

13. Pezzotti, N., Höllt, T., Lelieveldt, B., Eisemann, E. & Vilanova, A. Hierarchical Stochastic Neighbor Embedding. Comput. Graph. Forum 35, 21–30 (2016).

14. van Unen, V. et al. Mass Cytometry of the Human Mucosal Immune System Identifies Tissue- and Disease-Associated Immune Subsets. Immunity 44, 1227–1239 (2016).

15. van der Maaten, L. Accelerating t-SNE using Tree-Based Algorithms. J. Mach. Learn. Res. 3221–3245 (2014).

16. Pezzotti, N. et al. Approximated and User Steerable tSNE for Progressive Visual Analytics. IEEE Trans. Vis. Comput. Graph 23. 1739–1752 (2017).

17. Setty, M. et al. Wishbone identifies bifurcating developmental trajectories from single-cell data. Nat. Biotechnol. 34, 637–645 (2016).

18. Comaniciu, D. & Meer, P. Mean shift: a robust approach toward feature space analysis. IEEE Trans. Pattern Anal. Mach. Intell. 24, 603–619 (2002).

19. Spits, H. & Cupedo, T. Innate lymphoid cells: emerging insights in development, lineage relationships, and function. Annu. Rev. Immunol. 30, 647–675 (2012).

20. McKenzie, A. N. J., Spits, H. & Eberl, G. Innate Lymphoid Cells in Inflammation and Immunity. Immunity 41, 366–374 (2014).

21. Spits, H. et al. Innate lymphoid cells–a proposal for uniform nomenclature. Nat. Rev. Immunol. 13, 145–149 (2013).

22. Robinette, M. L. et al. Transcriptional programs define molecular characteristics of innate lymphoid cell classes and subsets. Nat. Immunol. 16, 306–317 (2015).

23. Schmitz, F. et al. Identification of a potential physiological precursor of aberrant cells in refractory coeliac disease type II. Gut 62, 509–519 (2013).

24. Schmitz, F. et al. The composition and differentiation potential of the duodenal intraepithelial innate lymphocyte compartment is altered in coeliac disease. Gut 65, 1269–1278 (2016).

25. Ettersperger, J. et al. Interleukin-15-DependentT-Cell-like Innate Intraepithelial Lymphocytes Develop in the Intestine and Transform into Lymphomas in Celiac Disease. Immunity 45, 610–625 (2016).

26. Mou, D., Espinosa, J., Lo, D. J. & Kirk, A. D. CD28 negative T cells: is their loss our gain? Am. J. Transplant 14, 2460–2466 (2014).

27. Bendall, S. C. et al. Single-cell mass cytometry of differential immune and drug responses across a human hematopoietic continuum. Science 332, 687–696 (2011).

28. Shaham, U. & Steinerberger, S. Stochastic Neighbor Embedding separates well-separated clusters. arXiv [stat.ML] (2017).

29. Höllt, T. et al. Cytosplore: Interactive Immune Cell Phenotyping for Large Single-Cell Datasets. Comput. Graph. Forum 35, 171–180 (2016).

30. Finck, R. et al. Normalization of mass cytometry data with bead standards. Cytometry A 83, 483–494 (2013).

